# STORM imaging buffer with refractive index matched to standard immersion oil

**DOI:** 10.1101/2023.03.07.531488

**Authors:** Youngseop Lee, Yeunho Lee, Minchol Lee, Donghoon Koo, Dongwoo Kim, Kangwon Lee, Jeongmin Kim

## Abstract

Stochastic optical reconstruction microscopy (STORM) provides exceptional super-resolution imaging by sparsely blinking individual dye molecules in thiol-containing media. STORM is now well-established for imaging thin biological specimens, and recent technological advancements have expanded its use to thick tissues. While the use of mounting media with an oil refractive index has been shown to reduce light scattering within tissues and thus greatly improve imaging depth and resolution in optical microscopy, the refractive index of STORM imaging buffers is typically water-like and oil-index (OI) buffers have never been considered for this purpose. In this study, we report a 3-pyridinemethanol-based STORM buffer that matches the refractive index of standard immersion oil. Our OI buffer exhibits similar superior performance in terms of photoswitching of Alexa Flour 647 dye and STORM image quality in fixed cells as conventional STORM buffers, despite having a completely different refractive index. Interestingly, it shows remarkable stability for at least 25 days, and potentially longer, which will enable STORM imaging of a large number of cells on a single prepared slide, as well as larger field-of-view imaging through multiple field stitching. By achieving perfect index matching with oil immersion objectives, OI buffers can produce accurate nanoscale morphology of thin biological specimens, without the need for complex microscope calibrations across sample depth. More importantly, our STORM buffer is expected to play a crucial role in lightsheet STORM applications for thick tissues by reducing light scattering, thereby leading to improved imaging depth and localization performance.

## INTRODUCTION

Stochastic optical reconstruction microscopy (STORM) [1] and direct STORM [2] provide superior imaging resolution down to 20 nm through localization of single molecule images of synthetic dyes. Thanks to its excellent super-resolution capabilities, STORM has been an important imaging tool to study the nanoscale morphology of cultured cells and thin tissue sections [3-5]. Typically, these thin specimens prepared for STORM imaging are excited using highly oblique illumination with a high numerical aperture (NA) oil immersion objective [6]. This leads to smaller point spread function (PSF) images of the fluorophore with minimal background fluorescence, which is optimal for higher localization precision [7]. However, such an intentionally introduced refractive index mismatch between the aqueous specimen and the immersion oil induces spherical aberration, reducing the achievable imaging depth to a few microns above the microscope coverslip. Therefore, it is not suitable for imaging thick samples like cell spheroids, tissues, and small animals, although important for more comprehensive biological studies [8, 9].

Several recent STORM methods have improved imaging depths to a few tens of microns. For example, adaptive optics was employed to correct optical aberrations due to the systematic refractive index mismatch between the aqueous sample and silicone oil as well as the index heterogeneity within the sample [10]. Lightsheet STORM approaches using water-based objectives provide confined illumination without systematic index mismatch, achieving single molecule imaging with high signal-to-background ratios [11-13]. However, for deeper STORM imaging in thick biological samples, it would be advantageous to use a sample mounting medium with a higher refractive index than water. This idea can be supported by the experimental results of multiphoton imaging of kidney tissue [14], where tissue scattering decreases significantly and the imaging depth increases from ∼10 μm to over 100 μm as the mounting index increases from 1.33 (water) to ∼1.51 (oil). Similarly, as shown in Figure S1, we observed a dramatic decrease in light scattering and background fluorescence in axial plane lightsheet imaging [15] of pollen grains when mounted with 2,2-thiodiethanol (TDE), whose refractive index is *n*_*D*_ = ∼1.516 at the *D* line (*λ* = ∼589 nm) [16]. We therefore anticipate that deep tissue STORM will perform best in terms of imaging depth and localization precision when using the STORM imaging buffer with a standard oil index.

Typical STORM imaging buffers used for photoswitching of synthetic dyes such as Alexa Fluor 647 (AF647) include a primary thiol (such as mercaptoethylamine (MEA) and beta-mercaptoethanol) and an enzymatic oxygen scavenging system (glucose, glucose oxidase and catalase; GLOX) in phosphate buffered saline (PBS) [17]. The imaging buffer is adjusted to a basic pH (∼8.0) and has a refractive index of *n*_*D*_ = ∼1.34, hereafter referred to as water-index (WI) buffer. The refractive index of WI buffer can be raised to 1.45 without degrading the photoswitching performance simply by adding 80% (v/v) glycerol or ∼60% (w/v) sucrose [18-20]. A commercial glycerol-based medium, Vectashield (*n* = 1.45), was also found to be suitable as a STORM imaging buffer for AF647 dye [21]. A STORM imaging buffer close to the oil index for AF647 was suggested with 75% (v/v) TDE in Vectashield, yielding *n*_*D*_ = 1.50 [21]. However, in our STORM experiments at an exact oil index (*n*_*D*_ = 1.515) with a mixture of 94:6 TDE-Vectashield, the single molecule density of AF647 was too low (<0.003 molecules/μm^2^) for STORM imaging even when activated with 405 nm light. In addition, Vectashield has been reported to cause fluorescence quenching of AF647 dye in direct STORM [22]. An alternative oil index buffer – mainly with 95% (v/v) TDE, 68 mM MEA, and GLOX – also showed a single molecule density of less than 0.015 molecules/μm^2^ for AF647, still too low for high-quality STORM imaging in our tests (Figure S2). Therefore, oil index STORM imaging buffers still need to be developed.

In this work, we report a STORM imaging buffer with the same refractive index as a standard immersion oil by adding 3-pyridinemethanol (3-PM) to the WI buffer formulation. We show that 3-PM has the optical properties that can be used as a sample mounting medium for fluorescence microscopy. Our oil-index (OI) buffer leads to good photoswitching of AF647 dyes suitable for STORM imaging and works stably for ∼3.6 weeks in sealed samples. For Alexa Fluor 555, the OI buffer exhibits better single molecule blinking than the standard WI buffer. We demonstrate three-dimensional (3D) STORM imaging of fixed COS-7 cells using OI buffer, providing image quality comparable to STORM imaging with conventional WI buffer.

## RESULTS

### Optical properties of 3-PM

We chose 3-PM because of its higher refractive index (*n*_*D*_ = 1.545) than the immersion oil and other favorable chemical properties, including good miscibility with water for refractive index tuning, near-neutral pH, and pH adjustability using HCl and NaOH if needed without precipitation. When dissolved in PBS, the refractive index of the mixture was linear for the 3-PM volume fraction, ranging from ∼1.33 (0% 3-PM) to 1.545 (100% 3-PM) as measured by a hand-held refractometer (PAL-RI, Atago) at room temperature (Figure 1a). Thus, by using ∼82% (v/v) 3-PM, a mounting buffer can be created with the same refractive index as the standard immersion oil (*n*_*D*_ = 1.515). This leaves ∼18% of the solution available for the separate preparation of aqueous components commonly used in typical STORM imaging buffer, such as MEA and GLOX. The 3-PM’s wide index adjustment range also covers various indices such as silicone oil (∼1.40) and glycerol (∼1.45) common for immersion objectives.

**Figure 1.**
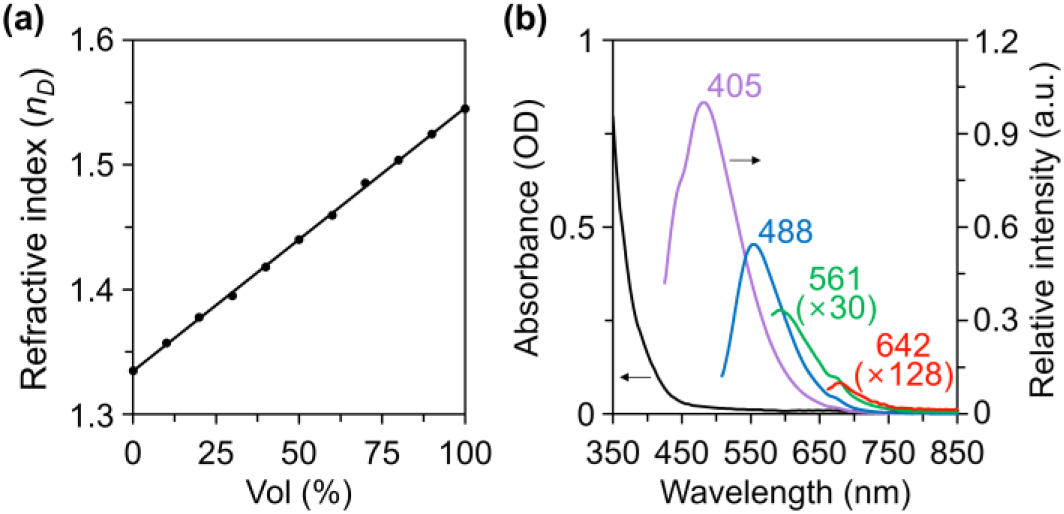
(a) Measured refractive index versus volume fraction of 3-PM in PBS. The linear fit is shown as a line. (b) Absorption (black) and emission spectra of pure 3-PM. Absorption length is 2.6 mm. The violet, blue, green, and red curves correspond to the emission spectrum when excited with 405, 488, 561, and 642 nm light, respectively. For clarity, the weak green and red emission spectra were scaled by 30× and 128×, respectively.

Pure 3-PM naturally exhibits clear light yellow color. To evaluate the optical transparency of 3-PM, we measured the optical absorption and emission spectra of pure 3-PM using a microplate reader (Synergy H1, BioTek) and a fluorescence spectrometer (FS-2, Scinco), respectively. As shown in Figure 1b, 3-PM appears to have relatively strong absorption at wavelengths below 450 nm, but the calculated absorption for a 100 μm propagation length is only 1.4% at the wavelength of 400 nm. Such absorption should be further reduced in OI buffer where 3-PM is diluted to 82%. Therefore, when used as a mounting medium for thick specimens such as tissue sections (typically, 30-100 μm in thickness), light absorption by 3-PM itself is negligible for all wavelengths from visible to near-infrared that are commonly used in microscopy. This is true for both the illumination light from the light source and the corresponding fluorescent light generated by the sample.

On the other hand, the emission of 3-PM when excited at the wavelengths of common laser lines (405, 488, 561, and 642 nm) exhibits a spectrum that partially overlaps the fluorescence wavelength range of organic dyes used in STORM (Figure 1b). The emission also becomes stronger when excited with shorter wavelengths of light. However, this emission was found to be much weaker than the typical fluorescence background of STORM. For example, under intense (on-axis) epi-illumination of 488 nm light at ∼4.7 kW/cm^2^ with weak 405 nm light added, the background signal of 82% 3-PM with a liquid thickness of ∼80 μm increased to less than 20 photons per pixel (130^2^ nm^2^ area for 20 ms) compared to deionized water. This background level is well below the typical 100-200 background photons per pixel in a typical 1.45-NA STORM system with on-axis epi-illumination. Indeed, a slight background increase was observed with Alexa Fluor 488 in single molecule experiments (Figure S3), but quality single molecule detection was still possible. We observed no noticeable background increase for Alexa Fluor 555 and 647 dyes. The 82% 3-PM can thus be considered a sufficiently transparent medium for use in conventional fluorescence microscopy and STORM.

### Photoswitching of Alexa Fluor 647 dyes in OI imaging buffer

As detailed in the Methods section, the OI imaging buffer at *n*_*D*_ = 1.515 was made by adding ∼82% (v/v) 3-PM while retaining the well-established composition of the conventional WI imaging buffer containing MEA and GLOX. We evaluated the OI imaging buffer by photoswitching Alexa Fluor 647, known as one of the best dyes for STORM. When excited by a 642 nm laser at 20 kW/cm^2^, AF647 dye dried on a coverslip and mounted in OI imaging buffer showed good single molecule switching (Figure 2a). The initial density of approximately 0.1 molecules/μm^2^ is well suited for STORM and is equivalent to that typically achieved with conventional WI imaging buffer at excitation intensities of ∼10 kW/cm^2^ (Figure 2b). As analyzed in Figure 2c, the initial density of single molecules in both OI and WI imaging buffers decreased over frames at similar rates.

**Figure 2.**
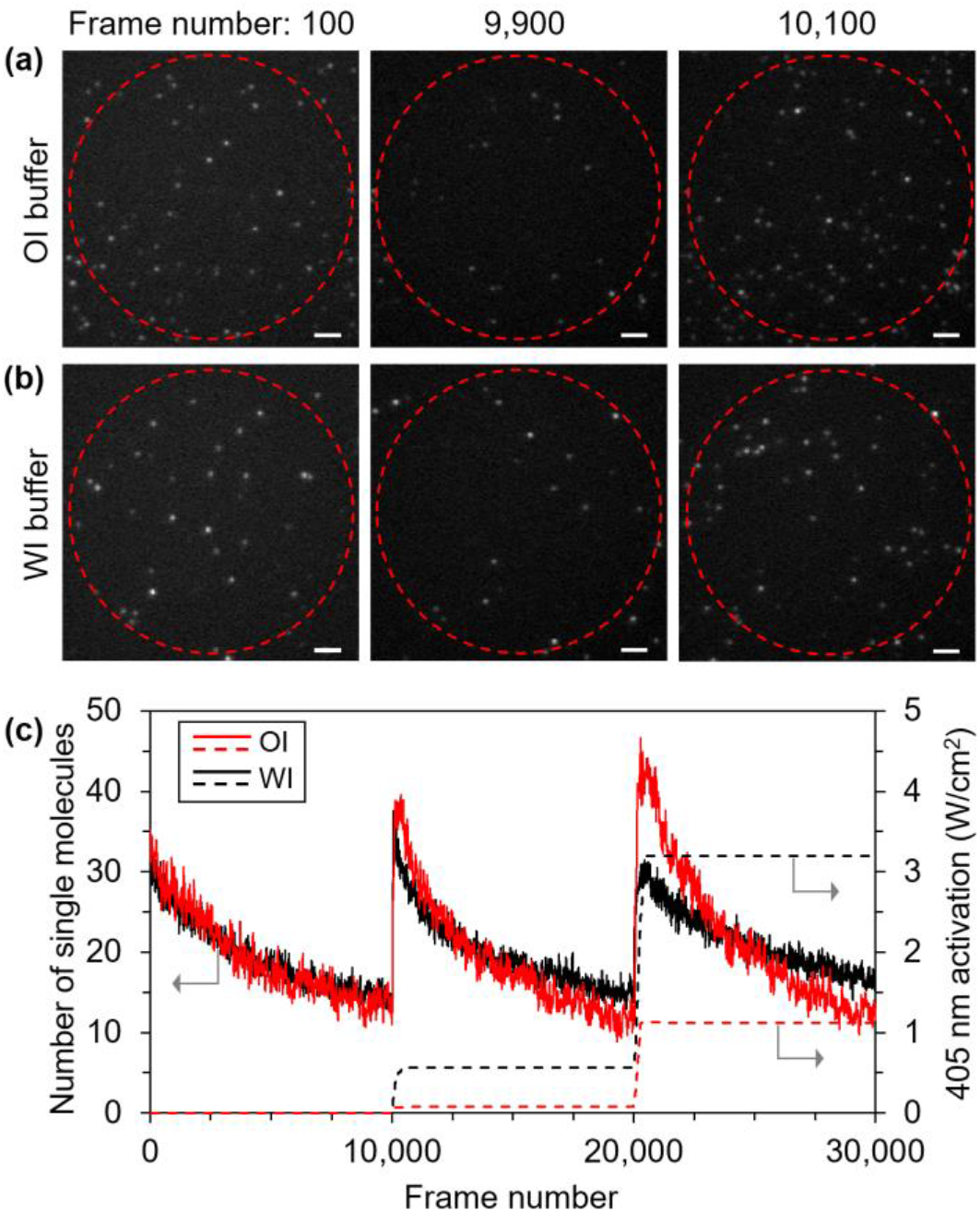
(a) Three representative image frames of Alexa Fluor 647 molecules in OI imaging buffer. (b) Control image frames of Alexa Fluor 647 molecules in WI imaging buffer. (c) The number of single molecules within the circular field of view (marked as red dashed lines in (a) and (b)) over image frames. Molecules appearing over several consecutive frames at the same location were counted as one molecule here. The optical activation using the 405 nm laser is indicated by dashed lines. Scale bars: 2 μm.

We then checked the optical activation of AF647 in OI imaging buffer. When weak illumination of 405 nm light was applied after about the 10,000^th^ and 20,000^th^ image frames, the single molecule density immediately recovered well to the initial population level, as shown in Figure 2. This recovery behavior is similar to AF647 mounted in WI imaging buffer, but interestingly, AF647 in OI imaging buffer exhibited 3-7 times higher sensitivity to activating light. From a practical point of view, the reduced intensity of the 405 nm activation light can help to minimize intrinsic autofluorescence [25], especially for thick biological specimens, which can lead to improved localization precision in STORM.

### Single molecule statistics of Alexa Fluor 647 dye

The photoswitching of AF647 dye in OI imaging buffer was further investigated at various 642 nm excitation intensities. It was found that AF647 molecules in OI imaging buffer turned off more slowly than in WI imaging buffer. More specifically, the median number of image frames over which dye molecules consecutively appear, defined here as “on-time”, was 1.2-1.9× longer in OI imaging buffer depending on illumination intensity (Figure 3a). During slower switching in OI buffer, AF647 molecules produced an average number of signal photons that were overall similar to dye molecules in WI buffer when each molecule’s signal photons were accumulated over “on-time” frames (Figure 3b). However, in contrast to WI imaging buffer which hardly changed the number of signal photons over illumination intensities of 10-30 kW/cm^2^ (also observed in Ref. [23]), OI buffer increased the number of signal photons with illumination intensity. The fluorescence background in both imaging buffers increased linearly with illumination intensity as expected, but OI buffer had a 10-20% higher background (Figure 3c). As a result, the average localization precision in OI imaging buffer was estimated to be nearly uniform at around 7.6 nm regardless of the excitation intensity (Figure 3d). This property differs from WI imaging buffer where the average localization precision of AF647 molecules is better at low excitation intensity within 10-30 kW/cm^2^, which was also demonstrated in Ref. [24].

**Figure 3.**
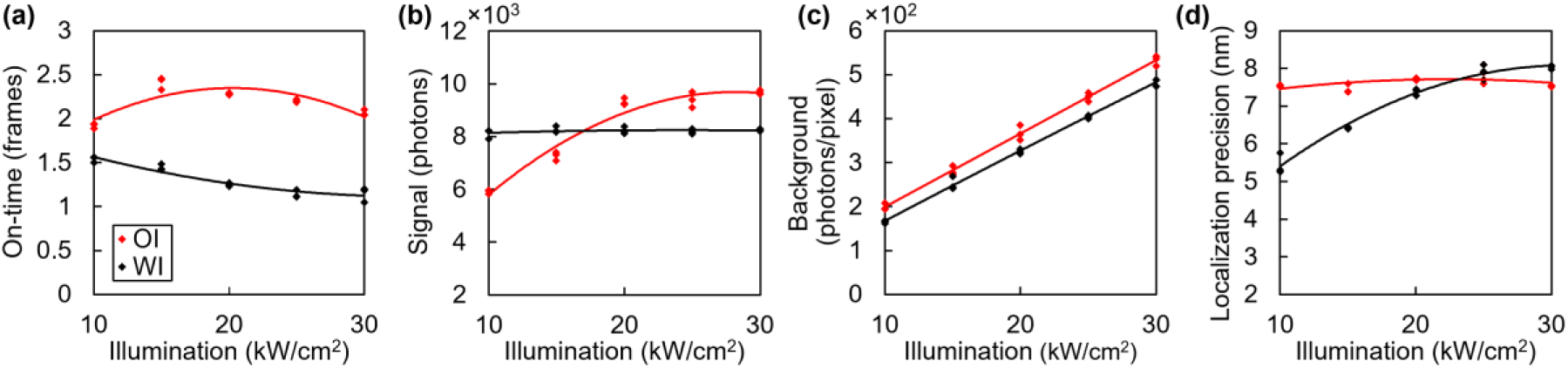
Single molecule statistics of AF647 dye in OI and WI imaging buffers versus illumination intensities. (a) Median of single molecule “on-time” distribution. (b) Average signal photons of single molecules. (c) Average background photons per pixel. (d) Average localization precision. Three different areas of the sample were imaged and analyzed for each illumination intensity, and the median/average value of the single molecule statistics of 30,000 image frames within each area was plotted as a dot. Lines are 2^nd^ order polynomial fits.

### STORM imaging of cells using OI imaging buffer

To confirm the biocompatibility of OI imaging buffer, we performed 3D STORM imaging of COS-7 cells using an astigmatic PSF through a cylindrical lens [25]. As shown in Figure 4, STORM images of AF647-labeled microtubules (α-tubulin) were of the same quality as microtubule images of cells mounted with conventional WI imaging buffer, when both were reconstructed from single molecule data of 40,000 image frames. In both OI and WI imaging buffers, individual microtubules appeared identical in a continuous manner with sufficient molecule density, which was not achieved with 40,000 frame data in TDE-based imaging buffer at oil index (Figure S2). Also, the tubular (or ring) structure of individual microtubules in OI imaging buffer was well resolved with a typical center-to-center distance of ∼31 nm in cross-sectional intensity profiles (Figure 4c), which was consistent with the results obtained from WI imaging buffer (Figure 4f) and is in good agreement with previously reported studies [24, 26].

**Figure 4.**
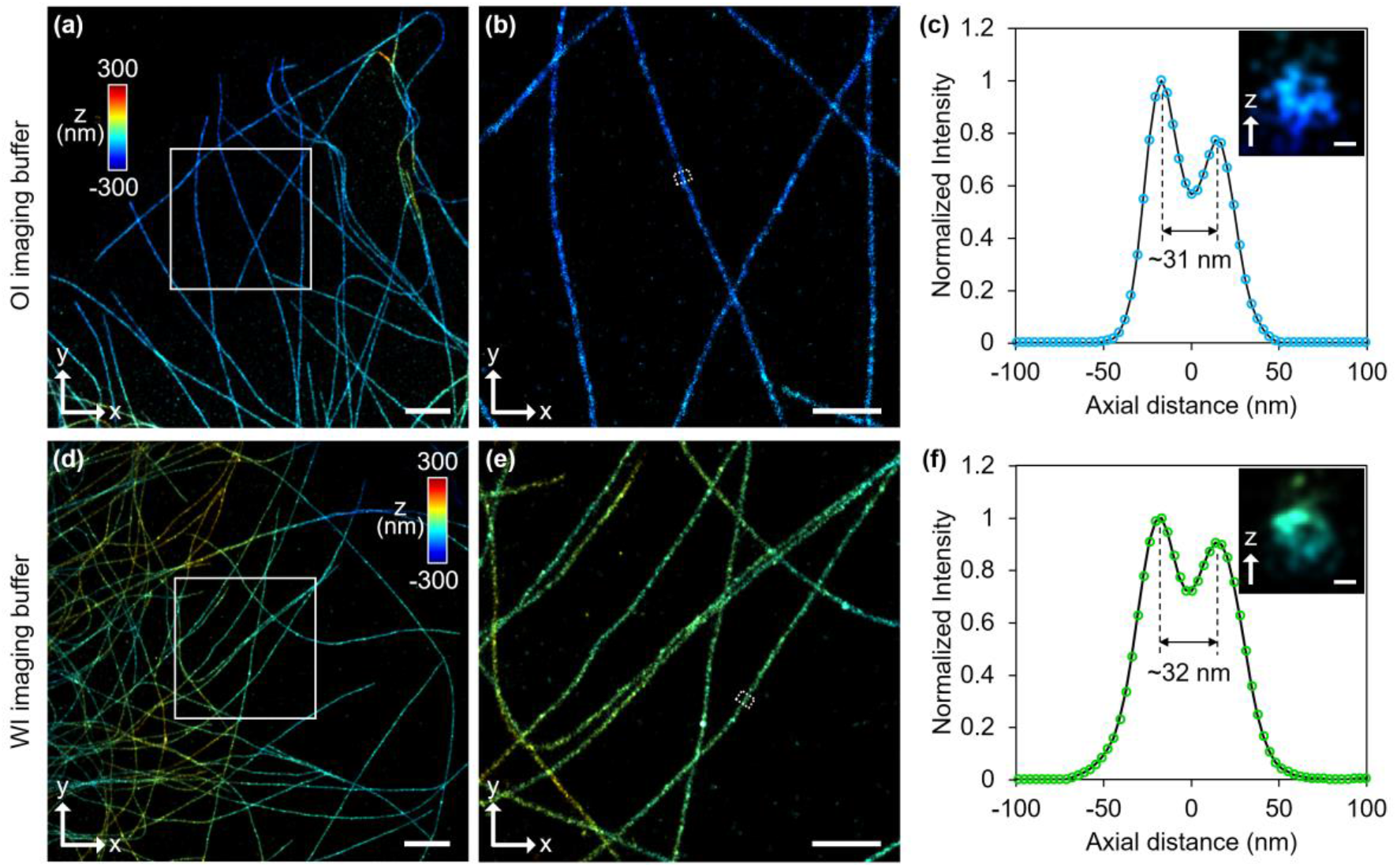
3D STORM imaging of AF647-labeled microtubules in COS-7 cells. (a, d) Super-resolution images of microtubule structures in OI and WI imaging buffers. (b, e) Magnified views of the boxed regions in (a) and (d), respectively. (c, f) Cross-sectional intensity profiles (projected in z) and images (insets) of the boxed areas in (b) and (e), respectively. Scale bars: 2 µm (a, d), 500 nm (b, e), and 20 nm (insets in c and f).

We also imaged the mitochondrial outer membrane (Tom20) immunostained with AF647 in COS-7 cells, another common target used to demonstrate STORM imaging. Mitochondrial networks were imaged well in OI imaging buffer and showed no noticeable difference from results in WI imaging buffer (Figure 5). Hollow-shaped membrane features, approximately 350-700 nm in diameter, were clearly resolved in cross-sectional views. These imaging results demonstrate that OI imaging buffer works well as a STORM imaging buffer for Alexa Fluor 647 dye and is fully compatible with fixed biological specimens. Moreover, our OI imaging buffer worked well for rhodamine-derived Alexa Fluor 555 (AF555), which does not blink well in the composition of WI imaging buffer for AF647 dye (Video 1). Indeed, the mitochondrial network labeled with AF555 was reconstructed as good as the network labeled with AF647, as shown in Figure S4. Finally, we evaluated the temporal stability of OI imaging buffer by performing daily STORM imaging on a COS-7 cell specimen as shown in Figure 6. After sealing the specimen with nail polish, we observed no signs of sample degradation in either conventional fluorescence images or STORM images of AF647-labeled microtubules for up to 25 days. Over the entire test period, the photoswitching behavior of AF647 dye remained similar and high-quality STORM images were also obtained. This is a distinct advantage compared to conventional WI imaging buffers, which last only a few hours after the sample is sealed due to GLOX-induced acidification [27, 28], as confirmed in the control experiment in Figure S5. The long-term stability of OI imaging buffer is probably due to its stable pH even when using GLOX. This could be attributed to the presence of pyridine derivates such as 3-PM, which are known to act as proton acceptors. An excess of 3-PM and its conjugated acid in OI imaging buffer could greatly enhance the buffering capacity, making the OI buffer insensitive to pH drops. We currently expect that specimens will continue to be stable for much longer than 25 days with longer testing. Even the ∼3.6 weeks tested is already approximately 100-150 times longer than the typical working time of conventional WI buffers, and long enough to obtain a large number of STORM images from a single specimen slide when needed.

**Figure 5.**
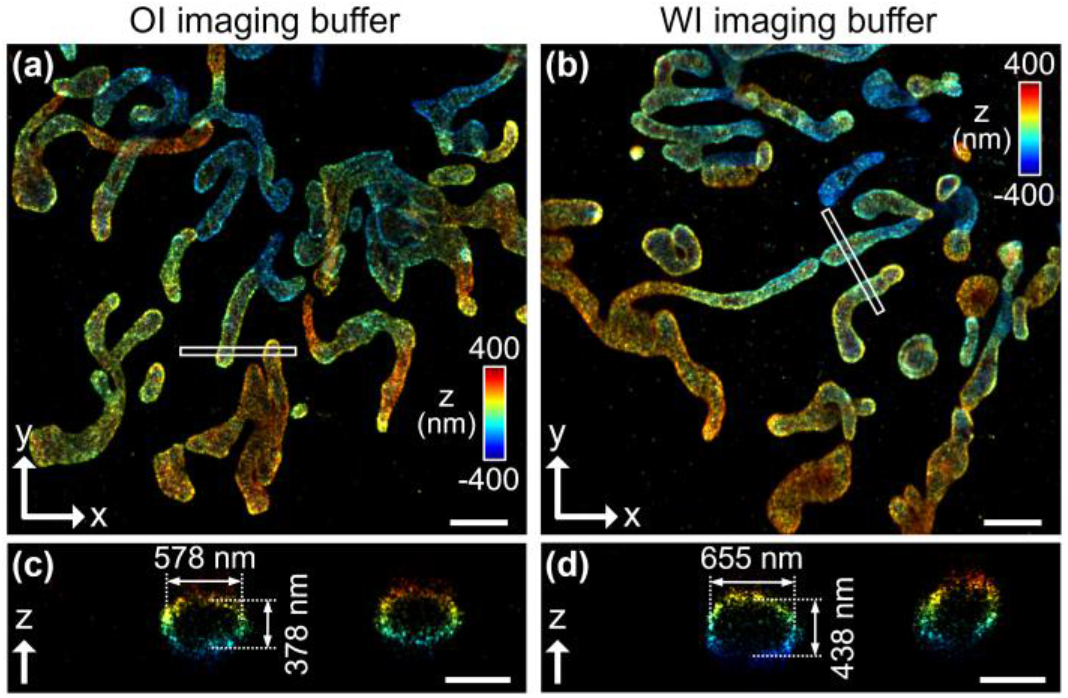
3D STORM images of AF647-labeled mitochondria network in COS-7 cells. (a, b) Super-resolution images of mitochondrial outer membranes in OI and WI imaging buffers reconstructed from 40,000 single molecule image frames. (c, d) Expanded z-plane views of the boxed regions in (a) and (b), respectively. Scale bars: 2 µm (a, b), 500 nm (c, d).

**Figure 6.**
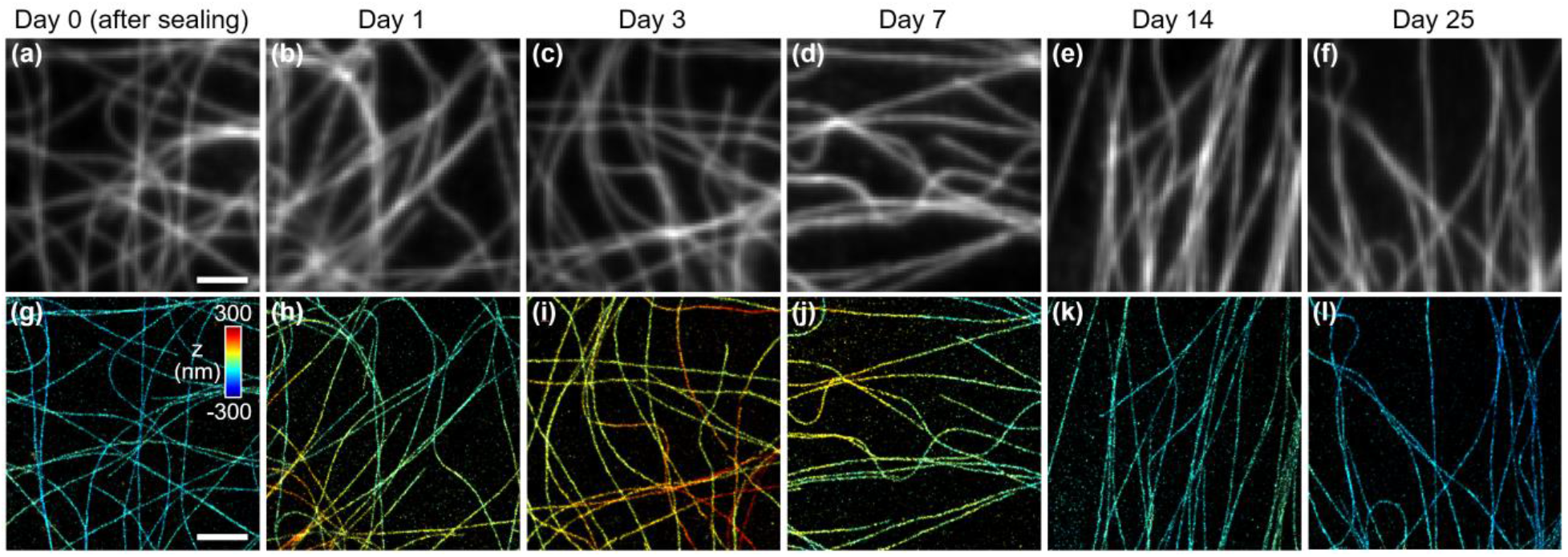
Temporal stability of a cell specimen when mounted with OI imaging buffer. (a-f) Conventional fluorescence images of AF647-labeled microtubules in COS-7 cells measured on different days. (g-l) Corresponding 3D STORM images. Scale bars: 2 µm.

## CONCLUSION AND DISCUSSION

For the first time, we have demonstrated a STORM imaging buffer whose refractive index perfectly matches standard immersion oil. The OI imaging buffer, which is based on 3-PM, is optically transparent and can effectively serve as a STORM mounting medium for Alexa Fluor 647 dye. Compared to the traditional WI imaging buffer, it has been proven to exhibit comparable performance in terms of single molecule switching and super-resolution cellular image quality. In terms of stability, the OI buffer in a sealed specimen has been tested to last for ∼3.6 weeks and may even last longer than that, even when the enzymatic oxygen scavenging system is in use. This is in stark contrast to the conventional WI buffer that can only be used for several hours due to pH drop. Therefore, OI imaging buffer is suitable for STORM imaging of a massive number of cells on a single prepared sample slide for large-scale analysis of biological processes. The observed long-term stability can also be useful for larger field-of-view STORM imaging through multi-field stitching in specimens such as sectioned tissues. Additionally, the OI imaging buffer works well for Alexa Fluor 555, making two-color STORM convenient as it can substitute for Cy3B, which is not available as an antibody conjugate.

The OI imaging buffer can be ideal for STORM applications using oil immersion objectives, because the buffer has a perfectly matched refractive index, which maintains the PSF shape along imaging depth without dimensional distortion. As a result, STORM imaging of thin specimens can be performed more accurately and conveniently without the need for complicated PSF calibrations based on field location and imaging depth [29, 30], provided the background fluorescence is weak. More importantly, the OI imaging buffer holds great potential for lightsheet STORM of thick biological specimens such as tissues and small animals, as it will lead to lower light scattering and a smaller and brighter PSF when compared to WI imaging buffer and water immersion objectives. This can thus improve localization precision from single molecule images at high signal-to-background ratios and also extend imaging depth with uniform through-depth imaging resolution.

The mechanisms behind the observed longer “on-time” of Alexa Fluor 647’s photoswitching and its higher sensitivity to 405 nm activation in OI imaging buffer are currently unclear and beyond the scope of this study. It is speculated, however, that 3-PM in the buffer hinders the binding between MEA and AF647 molecules and destabilizes the MEA-AF647 bonds, which may lead to each of the observed phenomena, respectively. Further investigation is required to fully understand this photochemistry. Besides Alexa Fluor 555 and 647 dyes, the compatibility of OI imaging buffer with other commercially available fluorophores is worth investigating. Future work may also include exploring modifications of OI imaging buffers with other thiols, alternative oxygen scavenging systems, and triplet state quenchers [28] for potential performance enhancement.

## MATERIAL AND METHODS

### The optical setup of STORM

The STORM setup was constructed based on a commercial microscope body (Eclipse Ti2-E, Nikon) on an optical table. For illumination, four lasers were used at wavelengths of 405 nm (50 mW, OBIS 405LX, Coherent), 488 nm (200 mW, Sapphire 488 HP, Coherent), 560 nm (1 W, 2RU-VFL-P-1000-560-B1R, MPB Communications), and 642 nm (1 W, 2RU-VFL-P-1000-642-B1R, MPB Communications). The laser beams were combined by three dichroic mirrors in free space, passed through an acousto-optic tunable filter (97-03151-01, G&H) for optical power control, and then coupled to a single-mode fiber (S405-XP, Thorlabs). The diverging laser light from the other end of the fiber was collimated and guided to the illumination port of the microscope using several folding mirrors. The collimated beam was focused on the back focal plane of the objective (100x/1.45, CFI Plan Apo Lambda, Nikon) through an achromatic lens (AC254-300-A, Thorlabs). The decenter of the incident beam and the achromatic lens to the illumination port was controlled by a motorized stage (PT1/M-Z8, Thorlabs) for off-axis illumination. For fluorescence imaging, we installed a quad-band dichroic beam splitter (ZT405/488/561/640rpcv2, Chroma) and an emission filter (ZET405/488/561/640m-TRFv2, Chroma) in the microscope filter cube and attached an electron multiplying charge-coupled device (EMCCD) camera (iXon Ultra 888, Andor Technology) to a customized camera port where a cylindrical lens (LJ1363RM-A, Thorlabs) can be inserted for 3D STORM. The z-stack images of orange (540/560 nm) and deep-red (633/660 nm) fluorescent beads (P7220, Invitrogen) mounted in OI or WI media were acquired and processed in SMAP [26] for 3D point spread function (PSF) calibration based on cubic spline PSF models for Alexa Fluor 555 and 647, respectively.

### Composition of STORM imaging buffer

WI imaging buffer (*n*_*D*_ = ∼1.341) consisted of 100 mM cysteamine (30070, Sigma-Aldrich), 10% (w/v) glucose (G7021, Sigma-Aldrich), 0.8 µg/ml glucose oxidase (G2133, Sigma-Aldrich), and 0.04 μg/ml catalase (C40, Sigma-Aldrich), 50 mM Tris-HCl (pH 8, Invitrogen), and 10 mM NaCl (S5150, Sigma-Aldrich) in phosphate-buffered saline (PBS). OI imaging buffer (*n*_*D*_ = ∼1.515) consisted of 100 mM cysteamine, 5% (w/v) glucose, 0.8 µg/ml glucose oxidase, 0.04 µg/ml catalase, 50 mM Tris-HCl, 3 mM NaCl, and 82% (v/v) 3-pyridinemethanol (A10381, Thermo Fisher Scientific) in PBS. To prepare the OI imaging buffer, we prepared three separate PBS-based solutions: (1) glucose with Tris-HCl and NaCl, (2) cysteamine with HCl for pH 8, and (3) glucose oxidase and catalase. These solutions were then mixed with 3-PM in a ratio of 11.4:6:1:82.6 to create the final OI imaging buffer. Alternative TDE imaging buffer (*n*_*D*_ = ∼1.515) tested consisted of 68 mM cysteamine (maximum solubility), 5% (w/v) glucose, 0.8 µg/ml glucose oxidase, 0.04 µg/ml catalase, 50 mM Tris-HCl, 3 mM NaCl, and 95% (v/v) TDE (166782, Sigma-Aldrich) in PBS. The pH of all three buffers was set to pH 7-8 with HCl.

### Sample preparation

*Alexa Fluor 647 dye on coverslips*: To evaluate the photoswitching property of fluorophores mounted in OI imaging buffer, AF647 secondary antibody (A31571, Invitrogen) was diluted to 1:10,000 in deionized water and dried on plasma-cleaned #1.5H coverslips (LH23.2, Carl Roth), followed by triple washes with deionized water. Dried samples were mounted on microscope glass slides with OI or WI imaging buffer and sealed immediately before imaging.

*Immunofluorescence of COS-7 cells*: Cells were cultured in T75 flasks according to standard culture protocols and seeded at ∼20% confluency on #1.5H coverslips (LH23.2, Carl Roth) immersed in wells of 12-well plates. When the cells had a stretched morphology after ∼2 days, they were fixed using a solution containing 3% paraformaldehyde and 0.5% glutaraldehyde in PBS and reduced using 0.1% sodium borohydride in PBS. Fixed cells were permeabilized with PBS with 0.5% Triton X-100 for 10 minutes and then blocked with blocking buffer (3% bovine serum albumin and 0.5% Triton X-100 in PBS). The cells were then incubated overnight at 4°C in primary antibody solutions with anti-α-tubulin (mouse monoclonal, 1:1,000, T9026, Sigma-Aldrich) or anti-TOMM20 (Rabbit polyclonal, 1:1,000, ab78547, Abcam) antibodies. After triple washes with washing buffer (10× diluted blocking buffer), cells were incubated for 45 minutes in secondary antibody solutions with AF647 secondary antibodies (anti-mouse, 1:1000, A31571, Invitrogen or anti-rabbit, 1:1000, A21245, Invitrogen) or Alexa Fluor 555 secondary antibodies (anti-rabbit, 1:1,000, A21428, Invitrogen), followed by triple washes with washing buffer and a final wash with PBS. Stained cells were mounted with OI or TB imaging buffer and sealed immediately prior to imaging. For OI imaging buffer, mounting was performed in four steps [16], sequentially applying three 3-PM solutions (10%, 25%, and 50% (v/v), respectively, in deionized water) and OI imaging buffer.

### Single molecule imaging experiment

*Imaging buffer evaluation*: Since AF647 dye on the coverslip was not uniformly distributed, conventional fluorescence images under weak 642 nm excitation showed field-to-field variation in the mean value of the image intensity histogram. For consistent photoswitching experiments, only imaging fields exhibiting similar mean intensities were selected as a way to match the initial population density of the fluorophores. At a chosen imaging area, we applied strong 642 nm excitation for ∼1 minute to turn off most of the fluorophores and then recorded 30,000 frames of single molecule images at 50 frames per second (fps). Weak 405 nm light (0-4 W/cm^2^) was used to examine optical activation sensitivity of AF647 dye. We used on-axis epi-illumination to ensure equal illumination intensities in experiments with OI and WI imaging buffers. For statistical analysis, 30,000 frame-long single molecule videos were repeatedly captured at different excitation intensities (10-30 kW/cm^2^) and different imaging areas of the sample while keeping single molecule density constant with 405 nm activating light. The intensity values for 642 nm illumination mentioned here are the average light power incident on the sample within the central field of view (diameter: ∼21 μm).

*3D STORM imaging of COS-7 cells*: We inserted the cylindrical lens used for 3D PSF calibration into the imaging beam path. After determining an imaging area of interest in the loaded cell sample with conventional fluorescence imaging mode, most AF647 molecules were turned off by intense laser excitation for 0-2 minutes. Single molecule videos were then acquired over 40,000 frames at 50 fps with the addition of 405 nm optical activation. The illumination intensity of the 642 nm laser light used for STORM imaging of microtubules and mitochondria outer membranes was ∼15 kW/cm^2^ and ∼10 kW/cm^2^ for the OI and WI imaging buffers, respectively. For imaging COS-7 cells labeled with Alexa Fluor 555, the illumination intensity of the 560 nm laser used was ∼8 kW/cm^2^ for both OI and WI imaging buffers and single molecule images of 40,000 frames were obtained at 50 fps with weak 405 nm activation. To access the long-term stability of OI imaging buffer, a COS-7 cell sample mounted with OI imaging buffer was imaged daily over a month. Each imaging experiment involved capturing a total of 40,000 single molecule image frames at 50 fps with 642 nm illumination (∼15 kW/cm^2^) and weak 405 nm activation. The sample was stored at 4°C between each imaging session.

### Single molecule data analysis and image reconstruction

The single molecule data obtained from buffer evaluation experiments were localized in two dimensions following the protocol described in Ref. [13], except that a circular Gaussian PSF model was used. Single molecule statistics was calculated only for molecules within the circular field of view with a diameter of ∼21 μm, corresponding to the FWHM of the Gaussian illumination beam profile. Single molecules appearing over consecutive image frames within 60 nm in diameter were regarded the same molecule, so signal photons were summed and background photons were averaged. The number of consecutive image frames over which individual dye molecule appears follows an exponential distribution, the median of which was defined as the “on-time” of the dye molecule. Localization precision was calculated analytically as derived from Ref. [31], assuming an excess noise factor of ∼2 in EMCCD cameras [32].

Single molecule videos for COS-7 cells were localized and rendered in SMAP software [26]. Single molecules with too weak (<1,000) or too strong (>10,000) signal photons were excluded during 3D image reconstruction. Axial (z) localization data for cells mounted in WI imaging buffer were corrected with a refractive index mismatch factor of 0.72 [18]. Sample drift was estimated and compensated by the correlation-based algorithm implemented in SMAP.

## Supporting Information

The Supporting Information is available free of charge. Autofluorescence imaging of pine pollen grains; STORM imaging of COS-7 cells mounted in TDE-based imaging buffer; Representative single molecule images of Alexa Fluor dyes; 3D STORM image of Alexa Fluor 555-labeled mitochondria in a COS-7 cell; Temporal stability of a cell specimen when mounted with conventional WI imaging buffer (PDF)

Representative single molecule images (200 frames at 50 frames/s) of Alexa Fluor 555-labeled COS7 cells when mounted with OI and WI buffers, respectively (Video 1) (MP4)

## Author Contributions

J.K. conceived the idea. Youngseop Lee (Y.L.^A^) and Yeunho Lee (Y.L.^B^) developed and tested imaging buffers. J.K. and M.L. built and calibrated the STORM setup. Y.L.^A^ prepared samples. Dongwoo Kim helped with cell preparation protocols and K.L. supported lab resources. Y.L.^A^ and Y.L.^B^ conducted imaging experiments. Y.L.^A^, Y.L.^B^ and Donghoon Koo analyzed STORM data. J.K. and Y.L.^A^ wrote the manuscript with input from all authors. J.K. supervised the research.

## Notes

The authors declare no competing financial interest.

## Acknowledgements

This work was supported by Creative-Pioneering Researchers Program through Seoul National University, the National Research Foundation of Korea (NRF) grant funded by the Korea government (MSIT) (No. 2020R1F1A1072519, No. 2021R1C1C1013067), and Korea Evaluation Institute of Industrial Technology (KEIT) grant funded by the Korea government (MOTIE) (No. 20018522). This work was in part supported by the Research Institute for Convergence Science. Donghoon Koo acknowledges a fellowship from the Hyundai Motor Chung Mong-Koo Foundation.

