## Supporting Information for "STORM imaging buffer with refractive index matched to standard immersion oil"

The Supporting Information contains five supplementry figures.

### Autofluorescence imaging of pine pollen grains

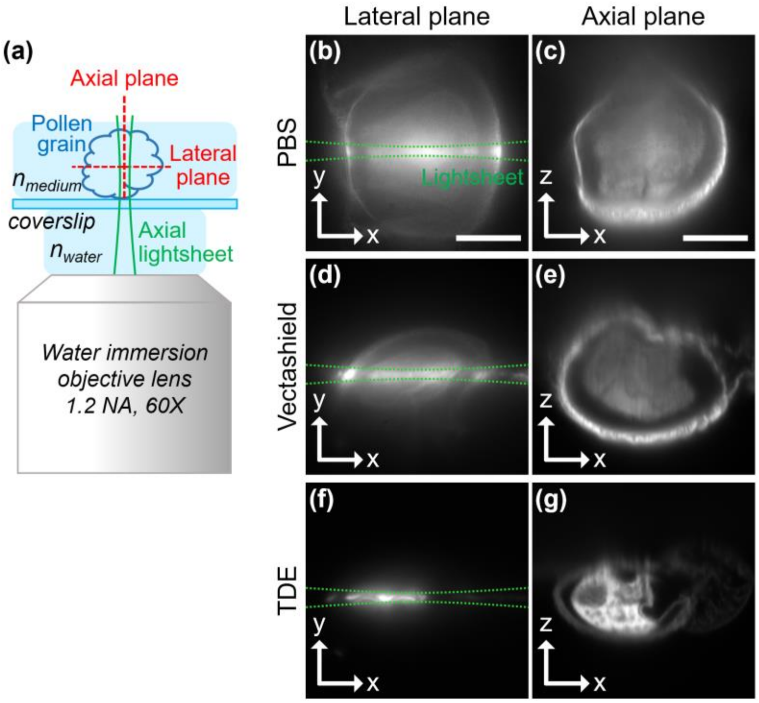


**Figure S1.** Autofluorescence imaging of pine pollen grains. (a) Schematic of axial plane lightsheet imaging. Pollen grains were excited by a 532 nm light sheet (FWHM thickness: ~1 μm at the beam waist) and the resulting autofluorescence images along the lateral and axial planes were captured. Lateral and axial plane images of pollen grains when mounted with (b, c) phosphate buffered saline (PBS; $n$ = 1.34), (d, e) Vectashield ($n$ = 1.45; H-1000, Vector Laboratories, Inc.), and (f, g) 2,2-thiodiethanol (TDE; $n$ = 1.52). The extent of lateral plane images is best limited within lightsheet illumination for pollen grains mounted at $n$ = 1.52, meaning minimal scattering of both excitation and emission light. The highest signal-to-background ratio of axial plane images is obtained when mounted at $n$ = 1.52. We note that, for $n$ = 1.45 or 1.52, axial plane images show spherical aberrations and axial dimensional distortion due to the use of a water immersion objective, but the conclusion that light scattering within the specimen is minimized when mounted with an oil index medium still remains valid. Scale bars: 20 μm.

### STORM imaging of COS-7 cells mounted in TDE-based imaging buffer

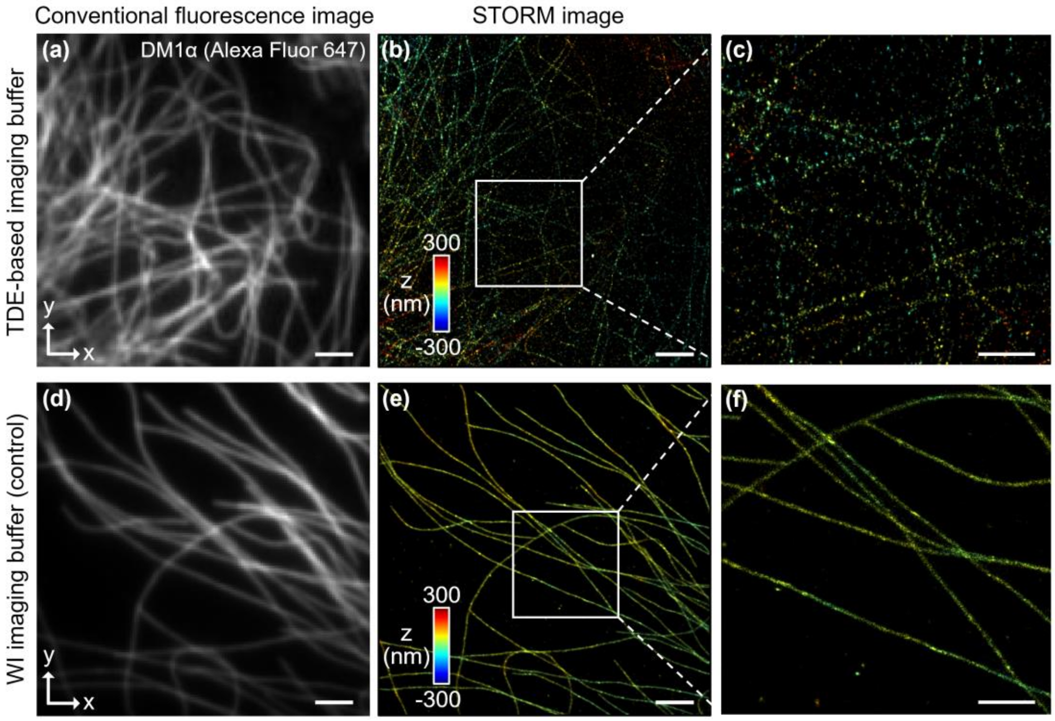


**Figure. S2.** STORM imaging of COS-7 cells mounted in TDE-based imaging buffer (95% TDE, 68 mM MEA; $n_D$ = 1.515). (a) Conventional fluorescence image of microtubules. (b) Corresponding 3D STORM image of (a). (c) Expanded view of the white boxed area in (b). Control experiment with conventional WI imaging buffer ($n_D$ = ~1.34): (d) Conventional fluorescence image. (e) Corresponding 3D STORM image of (d). (f) Expanded view of the white boxed area in (e). STORM images were reconstructed from single molecule data of 40,000 image frames acquired at 50 fps with an illumination intensity of 10 kW/cm$^2$. In TDE-based imaging buffer, the single molecule density is too low for high-quality STORM imaging even with continuous activation with 405 nm light. Scale bars: 2 μm (a, b, d, e) and 500 nm (c, f).

### Representative single molecule images of Alexa Fluor dyes

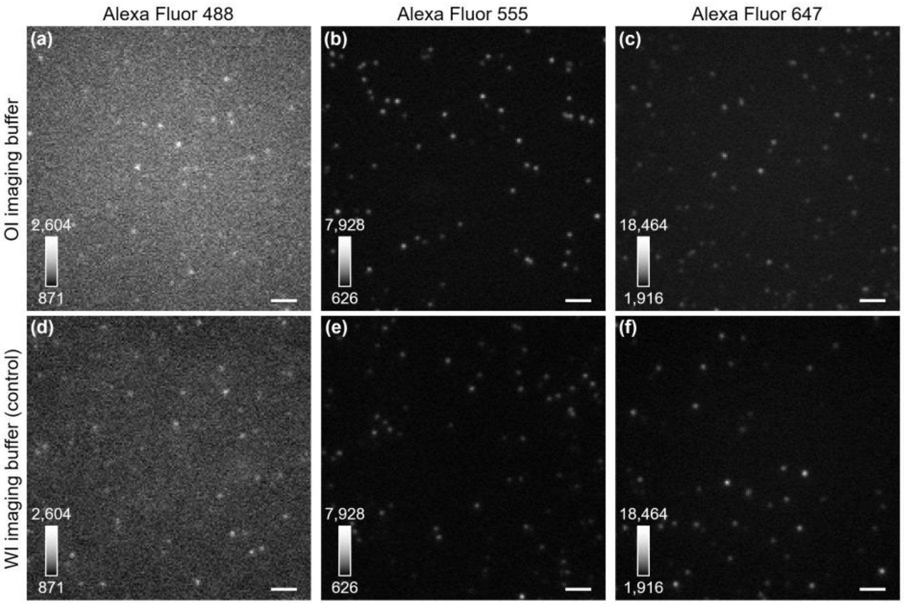


**Figure S3.** Representative single molecule images of Alexa Fluor dyes coated on coverslips. (a, d) Alexa Fluor 488 (AF488) dye mounted with OI and WI imaging buffers. Dye molecules were excited with a 488 nm laser with an illumination intensity of ~4.7 kW/cm$^2$. (b, e) Alexa Fluor 555 (AF555) dye mounted with OI and WI imaging buffers. Dye molecules were excited with a 560 nm laser with an illumination intensity of ~10 kW/cm$^2$. Imaging areas with low dye density were chosen as AF555 dyes in WI imaging buffer were not turned off well. (c, f) Alexa Fluor 647 (AF 647) dye mounted with OI and WI imaging buffers. Dye molecules were excited with a 642 nm laser with an illumination intensity of ~10 kW/cm$^2$. All images were taken at 50 fps. The image intensity is displayed as the 16-bit camera count, which includes a baseline of 500 counts. Image contrasts were set equally for each dye for direct comparison between different buffers. While AF488 dye showed a slightly higher background fluorescence in OI imaging buffer, single molecule imaging was still possible. There were no noticeable increases in background fluorescence in OI imaging buffers for AF555 and AF647 dyes. Scale bars: 2 μm.

### 3D STORM image of Alexa Fluor 555-labeled mitochondria in a COS-7 cell

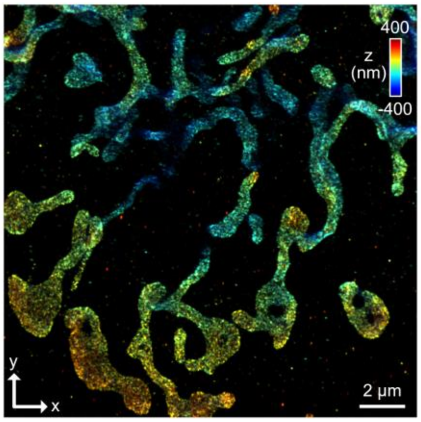


**Figure S4.** 3D STORM image of Alexa Fluor 555-labeled mitochondria in a COS-7 cell. The specimen was mounted with OI imaging buffer and 40,000 image frames of single molecule video were acquired for STORM.

### Temporal stability of a cell specimen when mounted with conventional WI imaging buffer

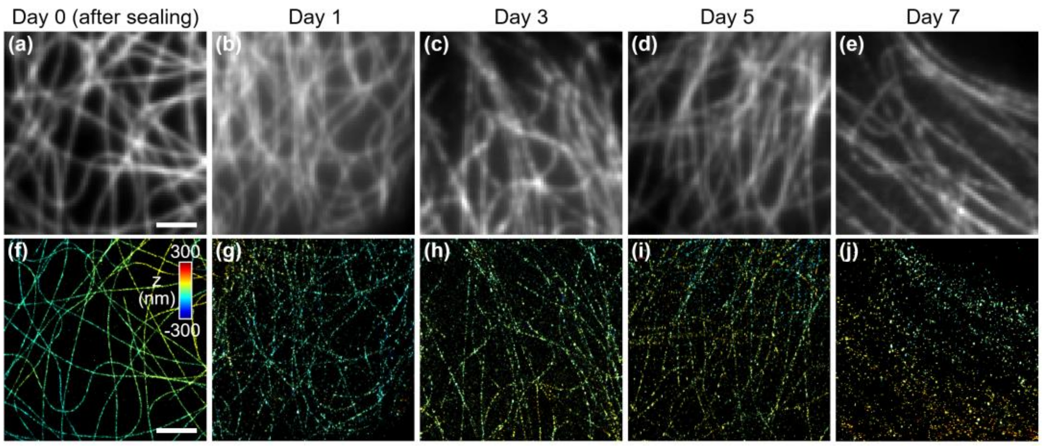


**Figure S5.** (a-e) Conventional fluorescence images of AF647-labeled microtubules in COS-7 cells measured on different days. There are a slight increase in background fluorescence and intensity fluctuations along each microtubule. (f-j) Corresponding 3D STORM images. High-quality STORM imaging is achieved only for several hours after the specimen is sealed. Scale bars: 2 μm.
